# Persistence of alternative reproductive tactics: a test of game theoretic predictions

**DOI:** 10.1101/2020.06.17.157727

**Authors:** Courtney R. Garrison, Scott K. Sakaluk, Ned A. Dochtermann

## Abstract

In many species, males produce signals to attract females. However, in some species and populations, only some males produce these signals with other males competing for and intercepting reproductive opportunities. In these systems, at least three tactics are expected: *always signal*, signal only when others are not (*assessors*), and *never signal*. The expected representation of these tactics within a population is frequently unknown in part because the costs of signaling (C) and the fitness value of a single reproductive bout (V) are difficult to quantify. Using a game-theoretic model, we predicted that the *always signal* strategy should only be present in a population if the fitness value of calling is greater than twice the cost (2C < V). We found that *always signal* males are apparently absent in decorated crickets (*Gryllodes sigillatus*), at least in our sampling of a laboratory housed population. Moreover, males were not strict assessors and instead signaled infrequently (30% of the time) when signaling by others was constant. Males also exhibited substantial among-individual variation in the propensity to call when other males were not signaling (τ = 0.3). Our results indicate a high relative cost of signaling (2C > V). The presence of among-individual variation in propensity to call is also suggestive of underlying genetic variation and a mixed evolutionary stable strategy. More generally, the apparent high cost of signaling and presence of variation in calling propensity suggests that reduced-cost strategies should spread quickly in populations.

## Introduction

The most dramatic behaviors of animals are often displays associated with attracting mates. Whether signaling genetic quality or condition (Kokko et al. 2003, Neff and Pitcher 2005), attainment of a specific threshold in quality (Takahashi et al. 2008), or parental ability (Hoelzer 1989), the importance of inter-sexual signaling has led to a diverse range of acoustic, visual, and chemical displays across animal taxa.

However, inter-sexual signals incur costs stemming from both their expression (e.g. energetic costs) and because they may attract predators or otherwise decrease the survival of signalers (White et al. 2022, Bernal et al. 2023). For example, parasitoids are attracted to the calls of cricket hosts (e.g. Gray et al. 2007), but these calls are necessary to reproduce. Individuals can avoid these costs via the expression of alternative reproductive tactics, such as by remaining silent and intercepting females orienting to signaling males (i.e. satellite individuals, Cade 1975, Cade 1981, Zuk et al. 2006). Indeed, tree crickets (*Oecanthus henryi*) switch from signaling to satellite when predation increases (Torsekar and Balakrishnan 2020). In some species and populations, these satellite individuals dramatically differ from signaling individuals phenotypically, even to the point of being physically incapable of producing signals (e.g. Zuk et al. 2006). In other species and populations, individuals can temporarily adopt either the satellite or signaling tactics despite heritability of calling propensity (Cade 1981).

The representation of these alternative reproductive tactics—the presence of multiple phenotypes associated with different routes to success in reproductive competition (Taborsky et al. 2008)—within a population can be predicted based on the relative benefits and costs of alternative tactics and modeled in terms of evolutionary games (Gross 1996, Walker and Cade 2003, Rotenberry et al. 2015). Under game theoretic models (Maynard-Smith 1982), populations are expected to reach equilibrium frequencies of signalers and satellites, which can include both tactics or only one. Here we develop a simple game-theoretic model of when individuals should signal or be satellites and use the predictions of this model to infer the <u>relative</u> costs and fitness benefits of signaling in the decorated cricket (*Gryllodes sigillatus*), a species in which some males adopt the satellite tactic (Sakaluk 1987). Because exact costs and fitness benefits can be difficult to quantify, this focus on the relationship between the two allows for clearer assessment of model predictions and will improve our general understanding of when alternative tactics are expected to be maintained in populations.

### A model of satellite behavior

In the case where males can plastically adopt either a satellite or signaling tactic, three general tactics might be expressed: *always signal, never signal*, and signal when no other males are signaling (i.e. *assess*). These tactics carry different costs (C) and fitness benefits (V) depending on which individual tactics are interacting. To understand what the relative costs and benefits of these tactics might be, we can start with an assumption that a non-signaling male can successfully intercept a female approaching a signaler from only one of four directions (Figure 1A). Thus, a *never signal* male interacting with either an *always signal* or *assess* male will have a payoff of 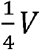, if we ignore additional costs to the satellite male, but a payoff of 0 when interacting with another *never signal* male (Figure 1B).

**Figure 1.**
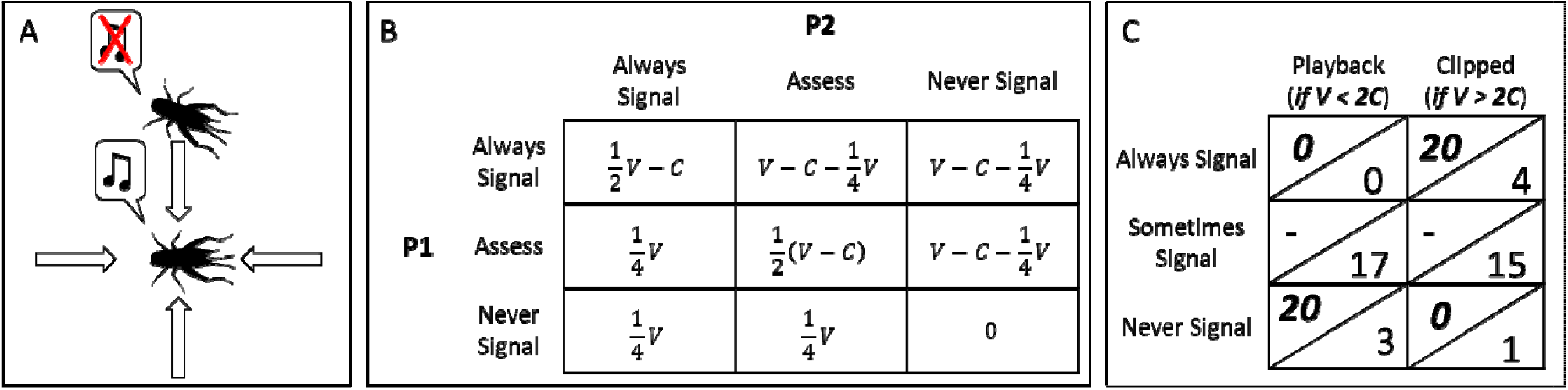
Assumptions, structure, analysis, and predictions from a game theory model of the interactions among individuals who *always signal, never signal*, and *assess*. A. Signaling males call (center cricket) and our model assumes females will approach from one of four directions (arrows). Satellite males (upper cricket) do not call and, instead, attempt to intercept females. We assume this satellite male will detect the female first if she approaches in the satellite male’s direction and therefore successfully mate with the female. (pictures courtesy of phylopic.org, Public Domain Mark 1 license) B. Reduced pay-off matrix for each of the tactics when played by individual P1 against another tactic played by P2. If V < 2C, the *always signal* tactic is not expected to persist in the population. If V > 2C, the *never signal* tactic is not expected to persist in the population. C. Predicted (above diagonal, bold) and observed (below diagonal) behavioral responses in the playback and clipped trials under two V:C relationships. The playback experiments directly test whether V<2C and the clipped wing experiments test whether V>2C. If V<2C, we expect no focal individuals to signal during the playback experiments. If V>2C, we expect focal individuals to always call while with a male who cannot signal. During playback trails no *always signal* individuals were detected, suggesting V < 2C. However, individuals did not strictly conform to the *always* or *never signal* tactics, instead sometimes signaling (middle row).

In natural settings we do not know what the exact payoff to a satellite might be: satellite tactics incur different costs than do signalers, including locomotive costs and changes in risk of predation (Raghuram et al. 2015, Torsekar et al. 2019, Geipel et al. 2020, Torsekar and Balakrishnan 2020). Individuals also can modify their calling based on more than just whether another individual is calling. For example, Blanchard’s cricket frog (*Acris crepitans blanchardi*) has been shown to lower dominant frequencies based on the properties of (synthetic) calls of other male frogs (Wagner 1989). Similarly, in other gryllid crickets, we previously found that indirect genetic effects affect multiple call components, indicating assessment when signaling (Garrison et al. 2020). Overall, the ability to detect, intercept, and successfully mate with the intercepted female will be dependent on a myriad of conditions that cannot be straightforwardly modeled. Nonetheless, both lower success and the different costs incurred by a satellite individual will be captured by the difference in payoff to satellite males and signaler. While we focus on the case when that relationship is 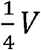 versus *V*, we also explore how changes in this affect our understanding of calling behavior in the discussion.

Further detailing the model, if two individuals with the tactic *always signal* interact, both will accrue the cost of producing signals but will have an equal probability of mating, leading to a pay-off of 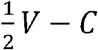 (Figure 1B). When an *always signal* male interacts with either an *assess or never signal* male, it incurs the cost of signal production (*C*) and has a 1 in 4 chance of losing a mating opportunity to a satellite opponent (Figure 1B). When two *assess* males interact, each has a probability of signaling half of the time, but when interacting with *always signal* males they have the *never signal* pay-off. The *assess* males also have the *always signal* pay-off when interacting with *never signal males*. These relationships can be extended across the possible combinations (Figure 1B). The full model description and analysis is available as supplemental material.

Analysis of this payoff matrix leads to several key predictions (Figure 1C, top left values in table cells): First, when fitness benefits (*V*) are <u>less</u> than twice the costs of calling (*C*), the *always signal* tactic is predicted to be lost from the population. Second, when fitness benefits are <u>more</u> than twice the costs, *never signal* individuals are lost from the population. Third, when benefits exceed four times the costs, only the *always signal* tactic persists. Fourth, when fitness benefits equal twice the costs of calling, all three tactics are expected to be maintained.

These general model predictions lead to specific testable predictions about the relative costs and benefits of signaling:

1. If a population contains *any* individuals that always signal, *V* ≥ *2*C.
2. If the population *only* contains individuals that always signal, then *V* > 4*C*.
3. If a population contains no always call males, instead consisting of satellite males that never call, then *V* ≤ 2*C*.

We tested for the presence of each tactic in *G. sigillatus* to allow an indirect assessment of *V* and *C*. We can test for the presence of *always signal* males by determining if a male ever calls while in an acoustic environment where there is constant signaling by other males. Similarly, we can test for the presence of never signal males by determining if a male never calls while in the presence of a non-calling male. Which tactics are present within a population is then informative as to the relative benefits and costs of signaling in this system.

## Methods

We tested the predictions of our model in banded crickets, *Gryllodes sigillatus*, from a sampled population descended from individuals captured in New Mexico (USA) and communally housed at high numbers (Ivy et al. 2005). These individuals had been isolated from predation and parasitoids and predators for ∼80 generations of random mating at the time of testing.

*G sigillatus* is ideal for testing questions regarding alternative reproductive tactics because males aggregate at high densities in natural settings (Sakaluk et al. 2002) and switch from signaling to satellite (S. Sakaluk, *pers. obs*.). We housed subjects individually in 0.71 L containers, provided *ad libitum* food (Purina Chick Starter) and water (provided in glass vials capped with cotton), and provided a shelter made of a small piece of cardboard egg carton. We moved crickets into individual housing when they were juveniles but all testing started after maturation. Testing occurred over a ∼two month period after maturation. This duration was necessary for the number of tests we sought to conduct (see below) but created the possibility of age-related changes in calling propensity (see Duffield et al. 2018). We maintained the cricket housing room, in which individuals were not acoustically isolated, at ∼ 27°C and on a 12:12 dark:light cycle reversed such that the room was dark during daytime hours, allowing us to conduct trials during the crickets’ “night”.

Two experiments were conducted to determine which tactics were present in the population. For all trials, individuals were moved to 0.71L containers that were then placed in Styrofoam chambers that provided acoustic isolation from neighboring chambers. These chambers were surrounded by 5 cm thick acoustic foam that provided additional sound insulation (Figure S1). During the clipped male tests (see below), we also added 5 cm acoustic foam over the front of the chambers to ensure calling by neighbors could not be heard. Prior observations demonstrated that calls of neighboring crickets were not detectable in the containers (Garrison 2017).

In the first experiment, playback tests, we observed 20 focal males during two-hour trials throughout which continuous signals of other male *G. sigillatus* were played. Playback was of single channel ∼3.5 minute audio file in which multiple calls were spliced together and then looped to allow continual playback. This file is available at https://github.com/DochtermannLab/CricketLoop. The playback speaker was placed ∼60 cm from the front of the acoustically isolated chambers and the front of the chambers were removed during these trials (Figure S1). Each focal individual was recorded during ten playback trials (separated by an average of 5 days). We recorded whether the focal male called during each trial. In this experiment *always signal* individuals would be expected to call during each trial. *Assess* and *never signal* individuals would be expected to not call during trials. If *V* < 2*C*, the above game-theory model would predict an absence of *always signal* individuals and that individuals adopting the *assess* and *never signal* tactics would not call (Figure 1C).

In the second experiment, clipped male tests, we observed the same 20 focal males as in the first experiment but while in the presence of non-focal males that had their wings surgically clipped following Stoffer and Walker (2012) and were thus unable to signal. Each focal individual was recorded during 10, two-hour trials. These trials were separated by an average of 5 days and were interspersed with the playback trials. To control for the possibility that the non-focal male identity influenced signaling propensity, all non-focal males were drawn from a population of highly inbred *G. sigillatus* (see Ivy et al. 2005 for details regarding the production of inbred individuals). We recorded whether the focal male called during each trial. Following the model described above, if *V* > 2*C*, there would be no *never signal* individuals and all individuals would call during the clipped-male bouts (Figure 1C).

Together, the playback and clipped-male testing scenarios resulted in 800 total hours of acoustic monitoring and allowed us to distinguish between *assess* individuals and always or never signal. These testing scenarios also allowed us to evaluate the relative magnitude of fitness benefits and costs of signaling. To test the predictions of our model we compared the predicted behaviors under different *V* vs. *C* conditions (Figure 1C) to the observed behavioral responses.

We also analyzed the outcomes of these trials using generalized linear mixed effects models. Whether an individual did not or did signal (0 versus 1) during a trial was analyzed as a binary variable with temperature (centered and scaled by its standard deviation), repetition number (1:10), mass (centered and scaled by its standard deviation), and time of day (centered and scaled by its standard deviation) included as fixed effects. Because of the length of testing, repetition number was included to capture the effects of habituation or age-related changes. A negative relationship between repetition number and propensity to call would indicate a decline in either motivation or capability. Individual ID was included as a random factor and among- and within-individual variances estimated, along with unadjusted repeatabilities (Nakagawa and Schielzeth 2010, Dingemanse and Dochtermann 2013). This statistical model was fit using the `lme4` package (Bates et al. 2007) in the R computing language. We also estimated the relative contribution of individual identity and the various fixed effects to behavioral variation using the `rptR` package (Stoffel et al. 2017).

## Results

### Calling propensity vis-à-vis game theoretic expectations

In the playback trials we predicted that, if *V* < 2*C*, there would be no individuals that always signaled (Figure 1C). Indeed, we found that no individuals always signaled. However, contrary to model assumptions, individuals in the playback trials did not wholly abstain from signaling, instead occasionally doing so (Figure 1C).

In the clipped-male trials, we predicted that there would be no individuals that never signaled if *V* > 2*C* (Figure 1C). In contrast, we did find that one individual—over 20 hours of recording—never signaled while in the presence of a clipped-male (Figure 1C). If *V* > 2*C*, we also predicted that all individuals would signal (i.e. both *assess* and *always call* individuals would call). Importantly, while males signaled most of the time during clipped-male trials (75% of bouts, Figure 2), individuals did refrain from signaling during some trials (Figure 1C).

**Figure 2.**
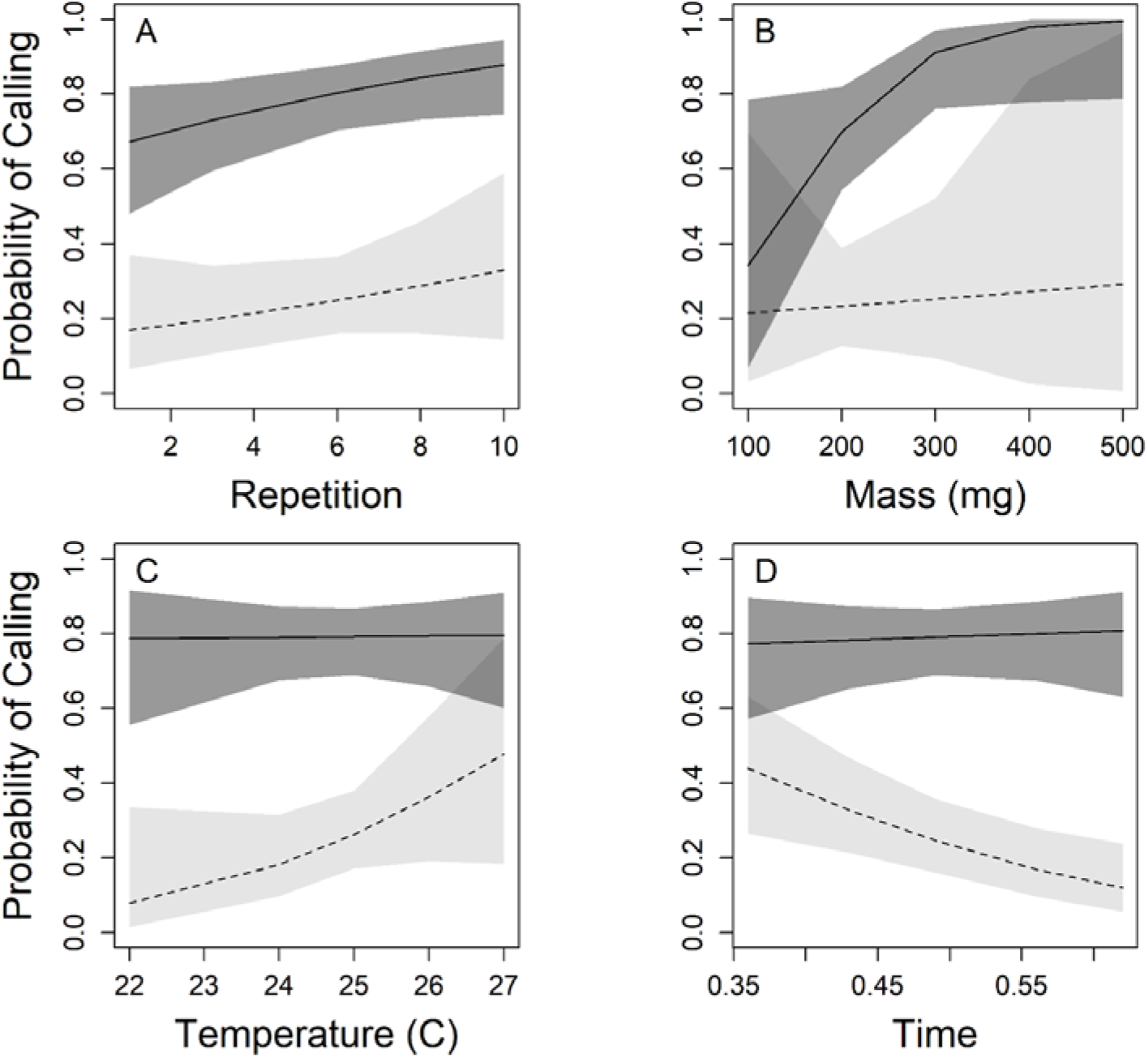
Relationship between probability of calling and (A) Repetition, i.e. over the sequence of recording bouts; (B) Mass, in miligrams; (C) Temperature, in degrees Celsius; and (D) Time of day (scaled from 0 to 1). In all panels the solid lines and dark gray shading correspond to clipped males. Dashed lines and light gray shading correspond to results from the playback experiment. Shading corresponds to 95% confidence intervals.

### General calling behavior

We found that crickets were significantly more likely to signal when in the presence of clipped males than during playback bouts (Figure 2, Table S1): males signaled during 74% of the clipped-male bouts, versus 35% of the playback bouts (Table S1; z = -3.07, p = 0.002). This is consistent with individuals adopting an assess tactic.

Probability of signaling did not significantly change across trials (Figure 2A, Table S1; z = 1.884, p = 0.06) or with temperature (Figure 2C, Table S1; z = 0.057, p = 0.95). The lack of a significant *negative* decline with repetition indicates that there were not age-related declines in calling that would have affected our inferences. Propensity to call increased with mass (Figure 2B, Table S1; z = 2.002, p = 0.045), a relationship that did not differ by treatment (Table S1). While there was no relationship between time and propensity to call when in the presence of clipped-males (Figure 2D, Table S1; z = 0.27, p = 0.79), during the playback bouts individuals decreased their propensity to call later into the dark period (Figure 2D, Table S1; z = -2.12, p = 0.034). Finally, we found that repeatability differed between the playback and clipped bouts. Repeatability during playback was 0.114 (p = 0.01) whereas it was 0.3 (p < 0.001) in the clipped-male bouts.

## Discussion

The lack of individuals who always signaled during the playback trials suggests the absence of *always call* individuals in our sampled population, though it is possible that they persist at a lower frequency than detectable here. The population, instead, being primarily composed of *assess* individuals that often, but not always, call indicates a mixed evolutionarily stable tactic. This is consistent with an individually based model of signaler-satellite dynamics that found support for populations with a mixed evolutionary stable strategy (Rotenberry et al. 2015). A mixed evolutionarily stable tactic implies that the fitness rewards of signaling are modest relative to costs.

Specifically, the combined results of the playback and clipped-male bouts are consistent with the fitness benefits of a single reproductive event being less than twice the costs of signaling (i.e. *V* < 2*C*). This is especially surprising given that this population has been free of selection from predation and parasitoids for ∼80 generations. Consequently, in this population, the costs of signaling were expected to only include metabolic costs.

If the crickets studied here indeed only incurred the energetic costs of calling, how large might these costs be? The energetic costs of calling in *G. sigillatus* are currently unknown, and these costs are highly variable across gryllid crickets: Hack (1998) found that the metabolic rate in house crickets (*Acheta domesticus*) while producing advertisement calls was approximately one and a half times an individual’s resting metabolic rate. In a similar range, Hoback and Wagner Jr (1997) estimated that advertisement calling corresponded to an increase of around two times the resting metabolic rate in variable field crickets (*Gryllus lineaticeps*). At the other extreme, calling mole crickets (*Gryllotalpa monaka*) increase their metabolism 13.5-fold relative to resting rates (White et al. 2008) and, for many crickets, calling requires similarly substantial increases in metabolic rates (e.g. Table 3 in Hack 1998). Based on these varied costs, it is unclear whether the relative costliness of calling in *G. sigillatus* is due to its energetic demands or low fitness benefits of calling.

The absence of *always call* individuals in our sample is also interesting independent of our model’s predictions given that calling effort has been shown to be condition-dependent in other gryllid crickets (e.g. Judge et al. 2008, Harrison et al. 2013, Duffield et al. 2018). Given that our crickets had *ad libitum* access to food, there should not have been condition imposed limits on calling effort. The absence of *always call* individuals is therefore unlikely to be tied to variation in condition. However, condition dependence in natural populations may result in some individuals expressing the *never call* strategy when unable to meet the costs necessitated by calling.

Regarding costs, it is also important to recognize that the satellite tactic might incur substantial and different costs than those incurred by signalers. Specifically, satellite males incur both higher energetic costs and changed predation risk due to moving to intercept females and around signalers. For example, female katydids—which, like crickets, actively move among males—experience higher predation from bats than do male katydids (Raghuram et al. 2015). This difference in predation is likely driven by a bias in bats toward scanning and approaching walking katydids versus still katydids (Geipel et al. 2020). However, costs are ecologically contingent: in a tree cricket, predation due to spiders is not sex-specific (Torsekar et al. 2019) and, in general, signaling increases mortality (White et al. 2022). Whatever the costs of a satellite tactic might be, they can nonetheless be represented within our existing model based on the relative pay-off to satellite individuals. Specifically, greater costs to satellites result in a decreasing fraction of *V*. Future modeling that explores changes in this fraction, with subsequent empirical testing, will reveal more about the relative costs satellites incur.

More generally, reduction in the probability of satellites intercepting females or increases in the costs faced by adopting the satellite tactic will increase the relative benefit of adopting calling. If the pay-off with a satellite tactic is indeed lower than we assumed, the absence of always call individuals here again becomes more surprising. This absence of males that always call despite higher costs for satellites would indicate even lower fitness benefits associated with signaling than already suggested by our results.

Comparing the number of individuals that called during either the playback or clipped-male trials to game theoretic expectations also demonstrates that individuals did not strictly conform to our model’s assumptions. For example, individuals did not conform strictly to either the assessor or never call strategies (Figure 1C). Whether this is because of insufficient complexity in our model or due to individuals assessing more cues than just the calling activity of other individuals is unclear at this point. If *assess* and *never call* represent a mixed evolutionarily stable tactic, this can be expressed either via all individuals expressing both tactics or by some individuals expressing one tactic and other individuals expressing the alternative. Individuals may also differ in underlying propensities to call, regardless of the acoustic environment they are in.

We were able to directly assess this possibility by repeatedly assaying individuals in each of the two environments (i.e. playback or silent). From this, we could estimate the degree to which individuals vary in their use of the assess or never call tactics. Here, we found evidence that only one individual expressed a pure tactic and all other individuals called during some trials but not during others. Estimated repeatabilities demonstrated the presence of among-individual variation in this propensity to call—at least when other males are not calling. Due to the relationship between repeatability and heritability (Boake 1989, Dingemanse and Dochtermann 2014, Dochtermann et al. 2015), this suggests genetic variation in the expression of alternative reproductive tactics. This connects to long-standing discussions about the distinction between tactics and strategies. Strategies and tactics are distinguished as strategies being the underlying propensity to express a particular phenotype, with tactics referring to that expressed phenotype ((Shuster and Wade 2003) but see (Taborsky et al. 2008) for criticisms of this distinction). Reversible and plastically expressed behaviors and morphologies represent tactics under this definition. If repeatability in the propensity to call is indicative of genetic variation in propensity, this would represent variation in underlying alternative reproductive strategies. Future research should determine both whether this among-individual variation is indeed indicative of genetic variation, as in other cricket species (Cade 1981), and if calling propensity is genetically correlated across acoustic environments. If so, genetic correlations may constrain the expression of optimal tactics.

More generally, our results have implications for expectations about the spread of alternative reproductive tactics or strategies in natural populations. If, as our results suggest, the relative fitness benefit of signaling is low, we would expect the rapid spread of non-calling tactics in populations. This will be particularly true when calling carries substantial costs. This prediction based on a low fitness benefit is consistent with the rapid spread of both satellite phenotypes and alternative signals in Hawaiian cricket populations after the introduction of parasitoids (Zuk et al. 2006, Tinghitella et al. 2018).

## Acknowledgements

We thank Tracie Ivy for the original collection of crickets and establishing inbred lines. We also thank R. Royauté and H. Sakaluk for important and useful discussions. This work was supported by US NSF IOS grant 1557951 to N.A.D. and NSF IOS grant 1654028 to S.K.S., Ben Sadd, and John Hunt.

## Ethical Note

All animals were treated humanely and adhered to ASAB/ABS Guidelines. This research was not reviewed by an institutional or governmental regulatory body because the species involved are not covered under applicable policies or statutes.

## Supplemental Materials

### S1. Model details and analysis

We developed a game theory model to determine the expected relative frequencies of *always signal, never signal*, and *assess* tactics within a population. Here we describe the model, starting with a description of the tactics that can be played and their pay-offs followed by the analysis of two tactic games and conclude with invasability analysis in the three tactic game. In these supplemental materials, we refer to behavioral phenotypes as tactics and strategies interchangeably. The use of “strategy” is more consistent with the literature on game-theory but is not used here to imply the presence or absence of total genetic control of a behavioral phenotype.

#### Playable tactics

Three tactics are available to be played: *always signal, never signal*, and *assess*. In a two person game each player can play any of the three tactics. Because each game is modeling a single male-male interaction and competition for a mate, pay-offs are determined by the fitness value of a single reproductive bout (*V*) and the costs of calling (*C*).

1. The never signal tactic models “satellite” behavior in which males attempt to intercept a female approaching a signaling male (Figure 1A). This tactic does not incur the cost of calling but cannot attract females in the absence of a signaling male. If an opponent male is signaling, we assume the never signal male can intercept a female approaching only from one of four directions (Figure 1A) and does so before the female can reach the signaling male. If two never signal tactics are played against each other, no female can be attracted and no fitness can be gained. Consequently, the average pay-off for this tactic against a calling male is 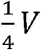 and 0 against another never signal.
2. The always signal tactic models males that signal regardless of which tactic is being played against. This tactic always incurs the cost, *C*, of signaling. If this tactic is played against another always signal tactic, each has a 50% chance of successful reproduction. If this tactic is played against a non-calling, satellite tactic, the calling player will lose one out of four reproductive opportunities but still incur the costs of signaling. The average pay-off for this tactic is therefore 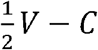 when played against another always signal tactic and 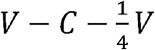 when played against non-signaling opponents.
3. The assess tactic models males who determine if opponents are signaling and do not signal if playing against always signal opponents. The assess tactic will, however, signal if playing against never signal opponents. When played against an always signal opponent, the assess tactic has the corresponding pay-off of the never signal tactic. When played against never signal opponents, the assess tactic has the pay-off of an always signal tactic. When assess tactics are played against each other, half the time the signal role is adopted and half the time the satellite role is adopted. The average pay-off is therefore: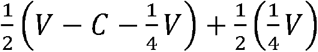 which expands to: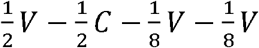 which simplifies to: 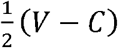.

Combined, these pay-offs can be summarized as (see also Figure 1B):

|  |  | P2 |  |  |
| --- | --- | --- | --- | --- |
|  |  | Always Signal | Assess | Never Signal |
| P1 | Always Signal | $\frac{1}{2}V - C, \frac{1}{2}V - C$ | $V - C - \frac{1}{4}V, \frac{1}{4}V$ | $V - C - \frac{1}{4}V, \frac{1}{4}V$ |
| | Assess | $\frac{1}{4}V, V - C - \frac{1}{4}V$ | $\frac{1}{2}(V - C), \frac{1}{2}(V - C)$ | $V - C - \frac{1}{4}V, \frac{1}{4}V$ |
| | Never Signal | $\frac{1}{4}V, V - C - \frac{1}{4}V$ | $\frac{1}{4}V, V - C - \frac{1}{4}V$ | 0, 0 |

#### A two-strategy game: Always Signal vs. Never Signal

We start by analyzing the case where the only available tactics are *always signal* and never signal. In this case, the pay-off matrix is:

|  |  | P2 |  |
| --- | --- | --- | --- |
|  |  | Always Signal | Never Signal |
| P1 | Always Signal | $\frac{1}{2}V - C, \frac{1}{2}V - C$ | $V - C - \frac{1}{4}V, \frac{1}{4}V$ |
| | Never Signal | $\frac{1}{4}V, V - C - \frac{1}{4}V$ | 0 |

Let *p* equal the frequency of always signal and 1 – *p* be the frequency of never signal. At equilibrium, mean pay-off of always signal equals mean pay-off of never signal. Thus:

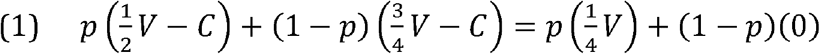

Expand terms:

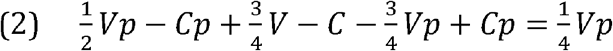

Simplifies to:

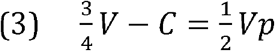

Solve for *p*:

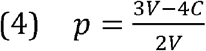

Given equation 4, for *always signal* to be evolutionarily stable and persist in a population 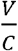 must be 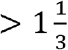. To get the 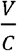 that results in a pure strategy of always signal, we set *p* to 1:

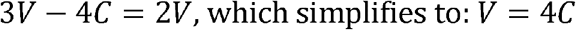

Consequently, whenever the benefits to signaling are four or more times the costs of signaling, *p* = 1 and the evolutionary stable strategy is a pure *always signal* strategy. When 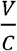 is between 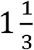 and 4, the evolutionary stable strategy will be a mixed strategy with the equilibrium frequency of the *always signal* and *never signal* strategies determined by equation 4. When V < 4/3, callers are lost, and alternative signaling modalities might be expected to prevail (e.g. cuticular hydrocarbons; Steiger et al. 2013).

#### Exploring the evolutionary stability of the three strategies

We explored the evolutionary stability of the three strategies by addressing the following questions: 1) is a population of *always signal* and never signal individuals at equilibrium vulnerable to invasion by an assess mutant; 2) is a pure population of *always signal* individuals, or a pure population of never signal individuals, vulnerable to invasion by an assess mutant; 3) is a pure population of assess vulnerable to invasion by either an *always signal* or never signal mutant. This was explored for different values of *V* and *C*.

1. We begin by considering an evolutionary stable population of caller and satellites in which *V*=20 and *C*=10 (i.e. the benefits of signaling to attract females is twice the cost of signal production). When *V* = *2C*, the always signal and never signal strategies have equal fitness. The equilibrium frequency of always signal individuals can be determined from equation 4 as above:

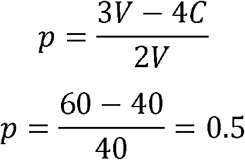

Thus, the frequency of always signal individuals is 0.5 and the never signal individuals is 1 – *p* or 0.5. The average payoffs with *V* = 20 and *C* = 10 are then:

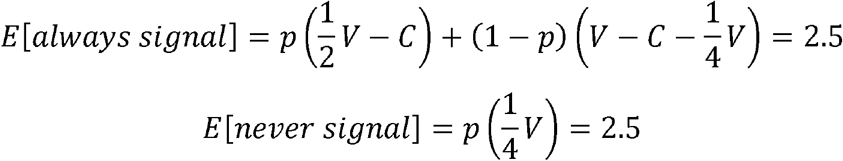
  1a. <u>Can this population be invaded by an assess mutant?</u> When *V* = *2C* half of the encounters involving the *assess* mutant will be with *always signal* individuals, and half will be with never signal individuals. The average payoff to the assess mutant is therefore:

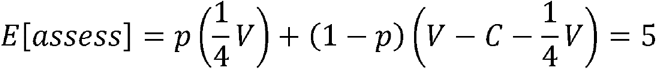

Thus, when *V = 2C*, assess can invade and spreads in the population.
  1b. <u>Can a mutant assess invade a population of pure caller?</u> Always signal individuals almost exclusively encounter other always signalers, whereas the mutant assess always encounters a signaler. Thus, the average payoff (i.e. fitness) of always signalers and assess when *V = 2C* is:

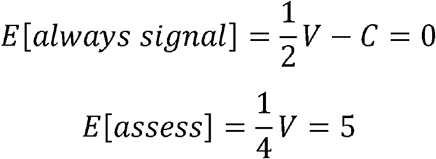

Thus, a pure always signal population is vulnerable to invasion by assess.
  1c. <u>Can a mutant assess invade a population of pure never signal?</u> Never signal individuals almost always encounter another never signaler, whereas the mutant assess always encounters a never signaler. The average payoff is therefore:

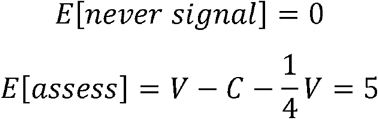

Thus, assess can invade and spread in a population of pure satellite.
  1d. <u>Can a population of pure assess be invaded by either a caller mutant or a satellite mutant?</u> Assess almost always encounters assess, and so too do the mutants. Thus, the average payoffs are:

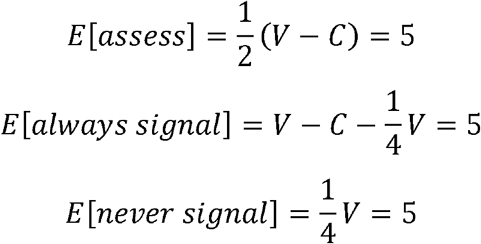

and at these values of *V* and *C* a pure assess population can be invaded by either a caller mutant or a satellite mutant but these mutants are not expected to spread, other than by drift.
2. We next examine when *V* ≠ *2C*. Consider the case where *V* = 30 and *C* = 10. In an evolutionary stable population of always signal and never signal the frequency of always signal (p) will be 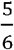, and the frequency of never signal will be 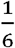 (equation 4). The corresponding payoffs are:

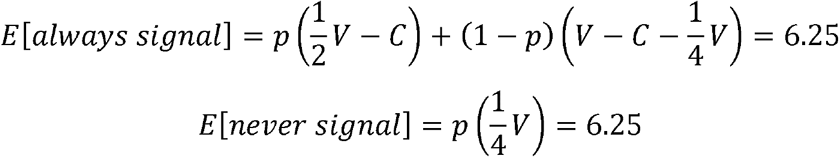
  2a. <u>Can this population be invaded by an assess mutant?</u> If *always signal* and never signal are in equilibrium, the payoff to assess is:

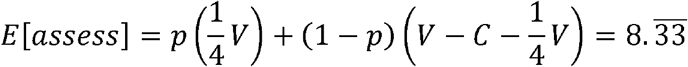

Under these conditions of V and C, assess can invade a population of callers and satellites at equilibrium and will spread.
  2b. <u>Assess can invade and spread in a population of pure callers:</u>

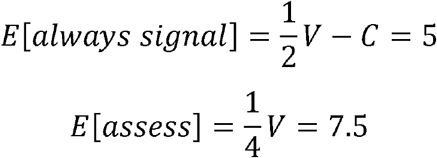
  2c. <u>Assess can also invade and spread in a population of pure satellites:</u>

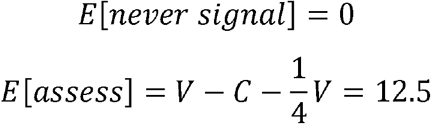
  2d. <u>Can a population of pure assess be invaded by either a caller mutant or a satellite mutant?</u>

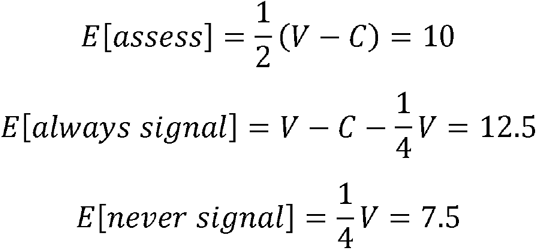

At these values of *V* and *C*, a pure assess population is vulnerable to invasion by an always signal mutant but cannot be invaded by the never signal strategy.
3. Now consider the case where *V* = 15 and *C* = 10. In a population of always signal and never signal at equilibrium the frequency of always signal (*p*) will be 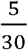, and the frequency of never signal will be .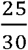. The corresponding payoffs are:

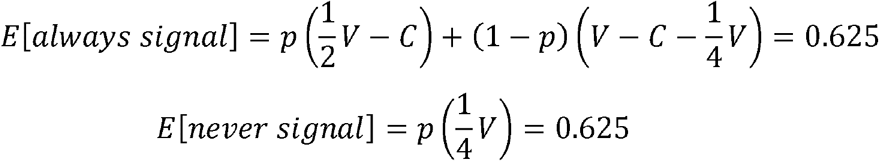
  3a. <u>Can this population be invaded by an assess mutant?</u> If always signal and never signal are in equilibrium, the payoff to assess is:

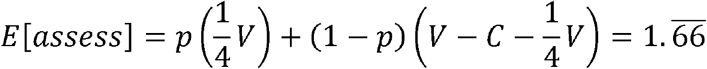

Under these conditions of *V* and *C*, assess can invade a population of callers and satellites at equilibrium and will spread.
  3b. <u>Assess can also invade a population of pure always signal:</u>

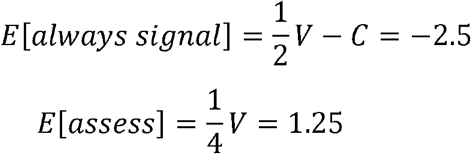
  3c. <u>Assess can invade a population of pure never signal:</u>

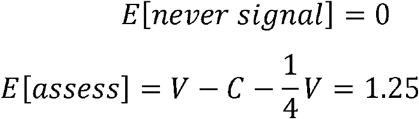
  3d. <u>Can a population of pure assess be invaded by either a caller mutant or a satellite mutant?</u>

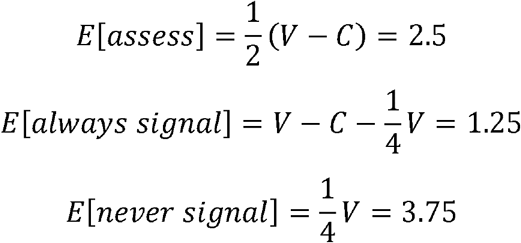

At these values of V and C, a pure assess population is vulnerable to invasion by a never signal mutant, but cannot be invaded by the always signal strategy.

#### Final thoughts and conclusions

The condition V = 2C is an important pivot point for the coexistence of the strategies. At these values of V and C, all three strategies confer the same fitness, and this seems to be the only condition under which all three strategies can coexist. This pivot point is almost certainly a function of the likelihood that a satellite male will succeed in intercepting a female orienting to a calling male. In our model, we assumed that a satellite male can only intercept females approaching from one of the four cardinal directions, and thus, the payoff to *never call* in this situation is V/4.

Although this is a reasonable starting approximation, it remains an open empirical question as to just how successful satellite males are at securing copulations with females orienting to calling males. Given that satellite males are likely to be able to intercept females from a greater range of directions but will not always be successful in courting these females, the true probability is unknown (and likely environmentally contingent). Consequently, it is worth noting that this assumption does not affect all predictions. Specifically, for a population to contain always call individuals, V must be greater than 2C (this is a necessary but not sufficient requirement). Our experiments allowed us to test this specific prediction.

When V >2C, some mixture of callers and assessors can be stable, whereas the pure satellite is excluded. When V<2C, some mixture of satellites and assessors is stable, whereas the pure caller is excluded.

Note that these conclusions appear to hold only when V/C lies between 1.33 and 4. When V/C >4, the ESS is pure caller, whereas when V/C <1.33, the ESS is pure satellite.

Empirically, this analysis could provide a powerful diagnostic tool for estimating costs and benefits of calling: whenever we find a population comprising satellites and males that behave something like assessors, it is likely that V < 2C; this is what seems to be the case for *Gryllodes sigillatus*.

---

**Table S1.**
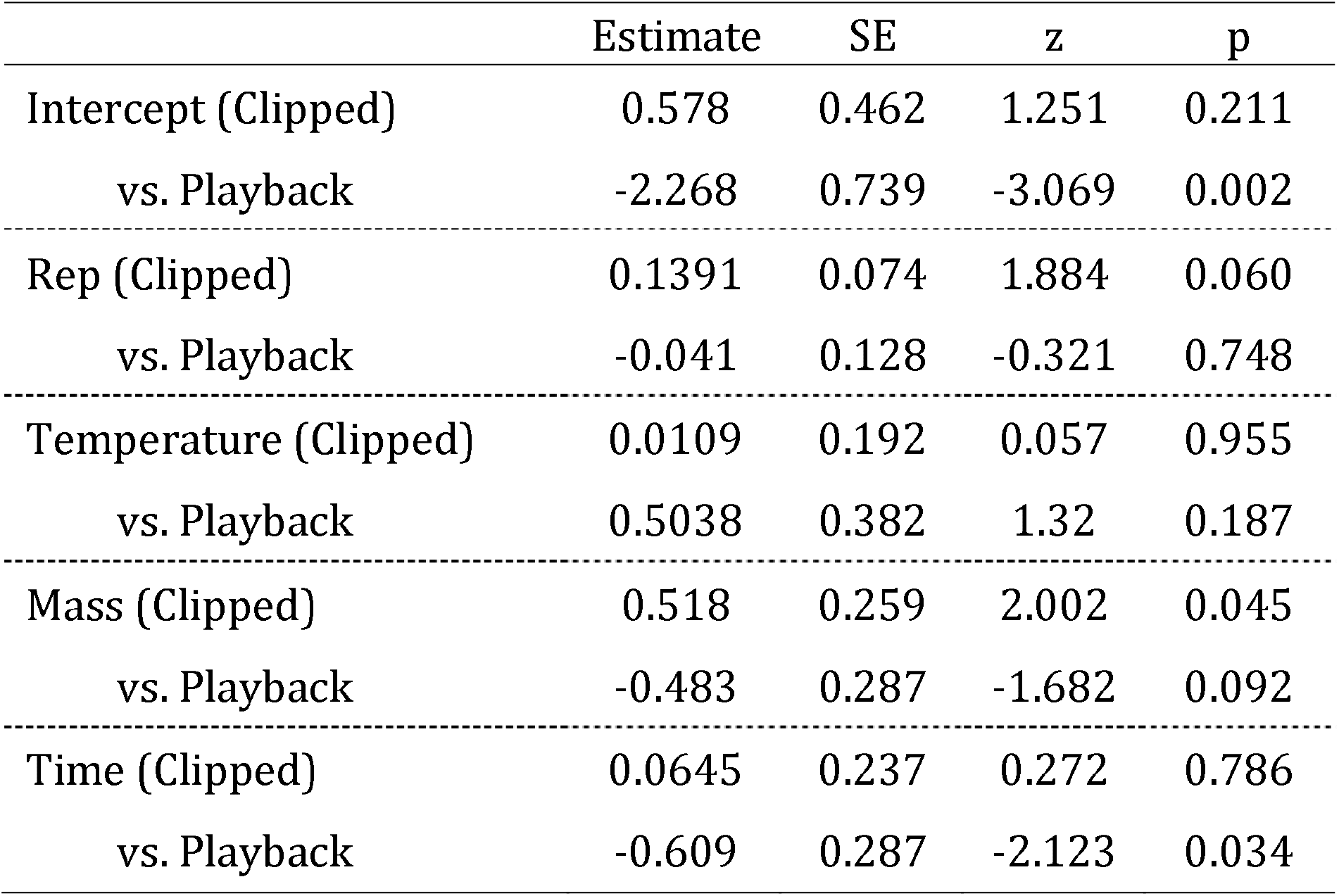
Summary of fixed effect estimates on behavioral responses. Results are from an analysis fit using the `glmer` function of the `lme4` package in R. The clipped treatment was fit as the intercept against which the estimate for the playback treatment was contrasted (i.e. the estimate for Playback would be the sum of the Clipped and Playback estimates). The effects repetition number temperature (Celsius, mean and standard deviation standardized) and mass (g, mean and standard deviation standardized) on calling propensity and interacted with treatment. Interaction estimates are presented as contrasts versus effects in the Clipped treatment. The model was fit with a binomial error structure.

**Figure S1.**
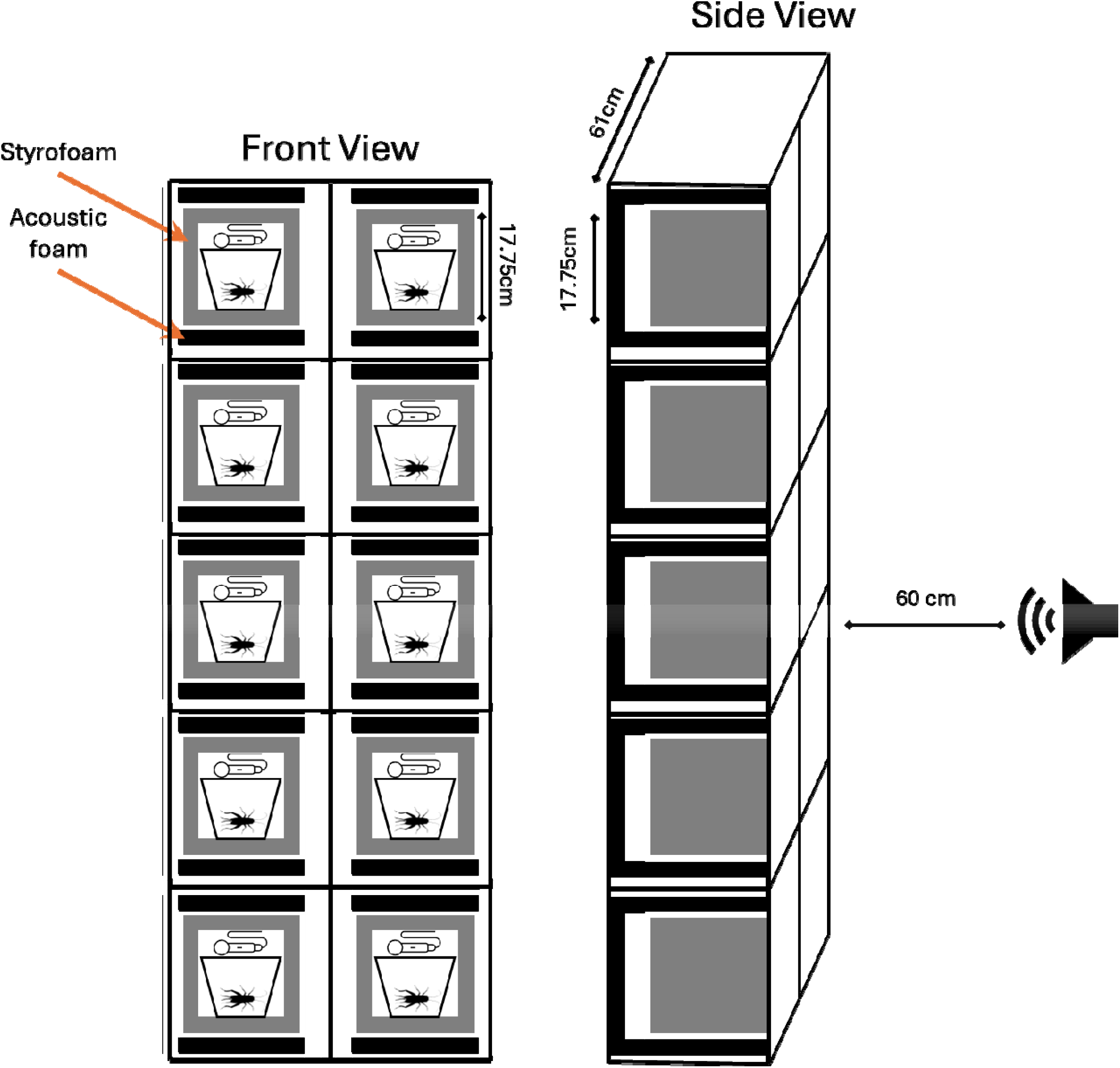
Schematic of recording chambers. During clipped-male trials 2.5 cm acoustic foam covers were placed over the front of chambers to provide acoustic isolation from neighboring chambers.

